# Partial alopecia area retains bulge hair follicle progenitor cells in Indian androgenetic alopecia patients

**DOI:** 10.1101/2023.09.06.556302

**Authors:** Akshay Hegde, Dyuti Saha, Savitha Somaiah, Colin Jamora

## Abstract

Androgenic alopecia (AGA) is a pathological condition characterized by the progressive decrease of scalp hair follicle density. Completely bald areas of AGA scalp still retain hair follicle stem cells but the progenitor cells are drastically decreased in number. However, it is unknown when the progenitor cells begin to diminish in number during the progression of AGA. Despite the prevalence of AGA in 58% of the Indian male population, no study to date has characterized the hair follicle stem and progenitor cell populations in AGA patients in this ethnic group. We observed that the partially bald area retains progenitor cells expressing CD34 and Sox9 but they are not present in the hair follicles in the completely bald area. Our study sheds light on the timeframe for effective therapeutic interventions based on modulating regulatory mechanisms to reinitiate the existing inactive follicle stem cells.

## Introduction

Androgenic alopecia (AGA), or male pattern hair loss, begins after the onset of puberty and progresses through adulthood (Ho et al., 2022). Studies report that AGA can affect 50% of men by the age of 50 and 50% of women by the age of 60 in the Caucasian population (Sinclair & Dawber, 2001). This can lead to decreased quality of life because of its significant impact on a person’s psychological well-being.

The hair follicle undergoes cyclical growth (anagen), regression (catagen) and resting (telogen) phases. In patients who are genetically predisposed for androgen-stimulated hair follicle miniaturization, the size of the scalp hair follicles are decreased with every cycle due to alterations in hair cycle dynamics. Eventually patients start producing microscopic hair due to the transformation of terminal hair to vellus (dormant) hair follicles (Kaliyadan et al., 2013). This results in the gradual replacement of large pigmented hair by depigmented hairs that are barely visible. Scalp skin biopsies from AGA patients exhibit decreased hair density, increase in the number of vellus hair, and altered hair follicle architecture (Sperling & Winton, 1990).

Hair follicle stem cells (HFSCs) reside in a niche in the hair follicle called the bulge. These HFSCs play a crucial role in the hair cycle by providing a continuous source of new cells for hair growth during the anagen phase. These stem cells are usually quiescent but during the anagen phase, they divide and give rise to transit amplifying progenitor cells (Purba et al., 2014), which are responsible for the growth of the hair shaft (Zhang et al., 2009). Studies suggest that therapies aimed at treating AGA can utilize either stem cells to reactivate hair follicles or regulatory mechanisms controlling hair follicle activity to stimulate hair growth (Talebzadeh & Talebzadeh, 2023). Realization of these therapeutic approaches are dependent upon understanding the status of the hair follicle stem and progenitor cells in AGA scalp. Previous reports have shown that in the bald area of the scalp in AGA patients, hair follicle stem cells are retained but progenitor cells are diminished indicating that the conversion of stem cells into progenitor cells is affected (Garza et al., 2011; Zhao & Hantash, 2014).

Recent studies conducted on Caucasian and Egyptian AGA patients have shown that, in the completely bald area of AGA scalp, HFSCs expressing K15 are present (Garza et al., 2011; Hend et al., 2015) but the progenitor cell population expressing CD200 and CD34 are diminished (Garza et al., 2011). This suggests that potentially AGA can be reversed by re-activating the conversion of HFSCs into progenitor cells (Mohammadi et al., 2016).

Previous studies have mainly focused on the fully bald area but hair follicles of the partially bald area of AGA have not been characterized for the status of stem and progenitor cells. Moreover, most of the studies have been conducted on Caucasian patients, although the prevalence of AGA differs with ethnicity (Ellis et al., 2002; Krupa et al., 2009; Wang et al., 2010). Though the prevalence is 58% in Indian males aged 30-50 years (Krupa et al., 2009), very few studies have been conducted on the Indian population. Thus in this study, we tested the presence of progenitor cells in the partially and fully bald scalp of Indian AGA patients.

## Results

### Bald area of AGA scalp lacks the complete structure of hair follicle and progenitor cells

The hair follicle is comprised of the infundibulum, bulge, sub-bulge, lower outer root sheath (ORS) and bulb based on the anatomy and the stem cell positions (Inoue et al., 2009a). To investigate the status of the hair follicle structure, we performed histological analysis of sections of hair follicles from the occipital area and bald frontal area of the AGA scalp (Fig 1A). This analysis revealed that the hair follicle from the occipital area contains the complete hair follicle structure whereas the bald frontal area lacks the bulge, sub-bulge, lower ORS and bulb structures. The remnant structure resembles the follicular ostia (Fig 1B). We further investigated the status of progenitor cells in these two regions from the AGA scalp. CD34^+^ progenitor cells are found in the outer root sheath (ORS) of the sub-bulge and suprabulbar region of the anagen hair follicle (Inoue et al., 2009b; Kloepper et al., 2008; Poblet et al., 2006). Another important population of cells expressing Sox9 are found in the ORS of the bulge, sub-bulge and proximal bulb area in the anagen stage (Purba et al., 2015). Immunostaining for these markers revealed that occipital unaffected area contains CD34^+^ and Sox9^+^ progenitor cells, whereas the completely bald area lacks them (Fig 1C and 1D). Our results are consistent with previous reports that the hair follicles of the bald area of AGA scalp from Caucasian patients lack progenitor cells derived from HFSCs (Garza et al., 2011) and also suggests that a cause of alopecia is due to a defect in the conversion of HFSCs to progenitor cells (Garza et al., 2011; Zhao & Hantash, 2014).

**Figure 1.**
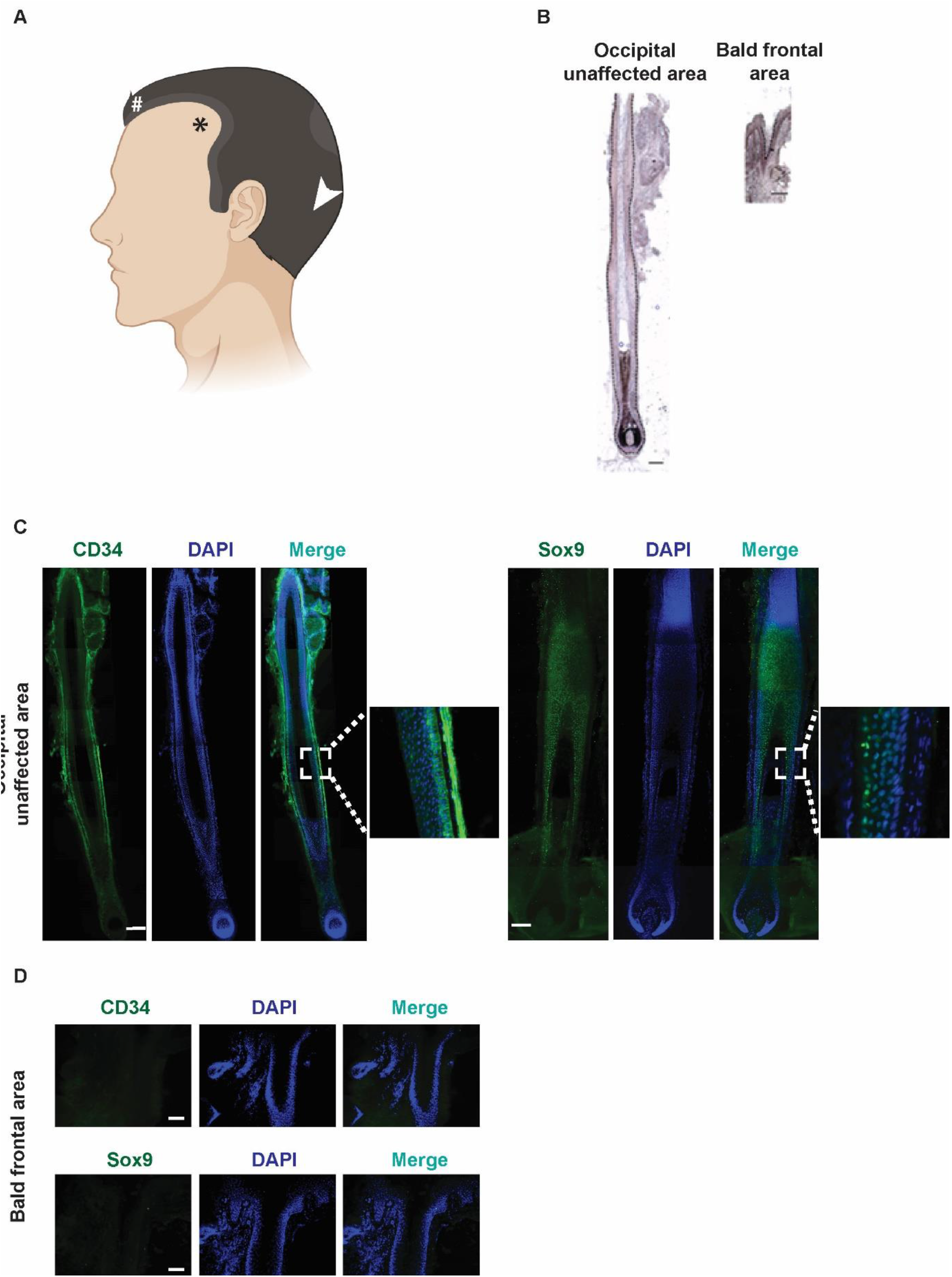
Bald frontal area of the AGA scalp lacks the complete structure of the hair follicle with absence of hair follicle progenitor cells. (A) Schematic of the scalp of an AGA patient marking the sites of hair follicle biopsy collection: Bald frontal area with vellus hair (marked by asterisk); Partially bald frontal area characterized by thinning/ recession (marked by hash); Occipital unaffected area with terminal hair (marked by arrowhead). (B) H&E staining of hair follicles from occipital unaffected area and the bald frontal area of the scalp. Immunofluorescence staining (green) for hair follicle progenitor cells marked by CD34 and Sox9 in (C) occipital unaffected area and (D) bald frontal area, respectively. Nuclei are marked by DAPI in blue. Scale bars: 100 µm. Images in (C) are composite images created by tiling several images together.

### Partially bald area of AGA scalp retains Sox9^+^ and CD34^+^ progenitors

An outstanding question in the field is when progenitor cells start to diminish. Thus, we tested the status of the progenitor cells in the partially bald frontal area of the AGA scalp. Immunostaining for CD34 and Sox9 markers revealed that the hair follicles of bald frontal area of AGA scalp lack the progenitor cells (Fig 1C) but they are still retained in the hair follicles of the partially bald frontal area (Fig 2A and 2B). Quantification of CD34^+^ and Sox9^+^ cells revealed that the number of progenitor cells are not significantly reduced in the partially bald area compared to the hair follicles of the occipital unaffected area. This suggests that loss of progenitor cells occurs after the partially bald stage of AGA.

**Figure 2.**
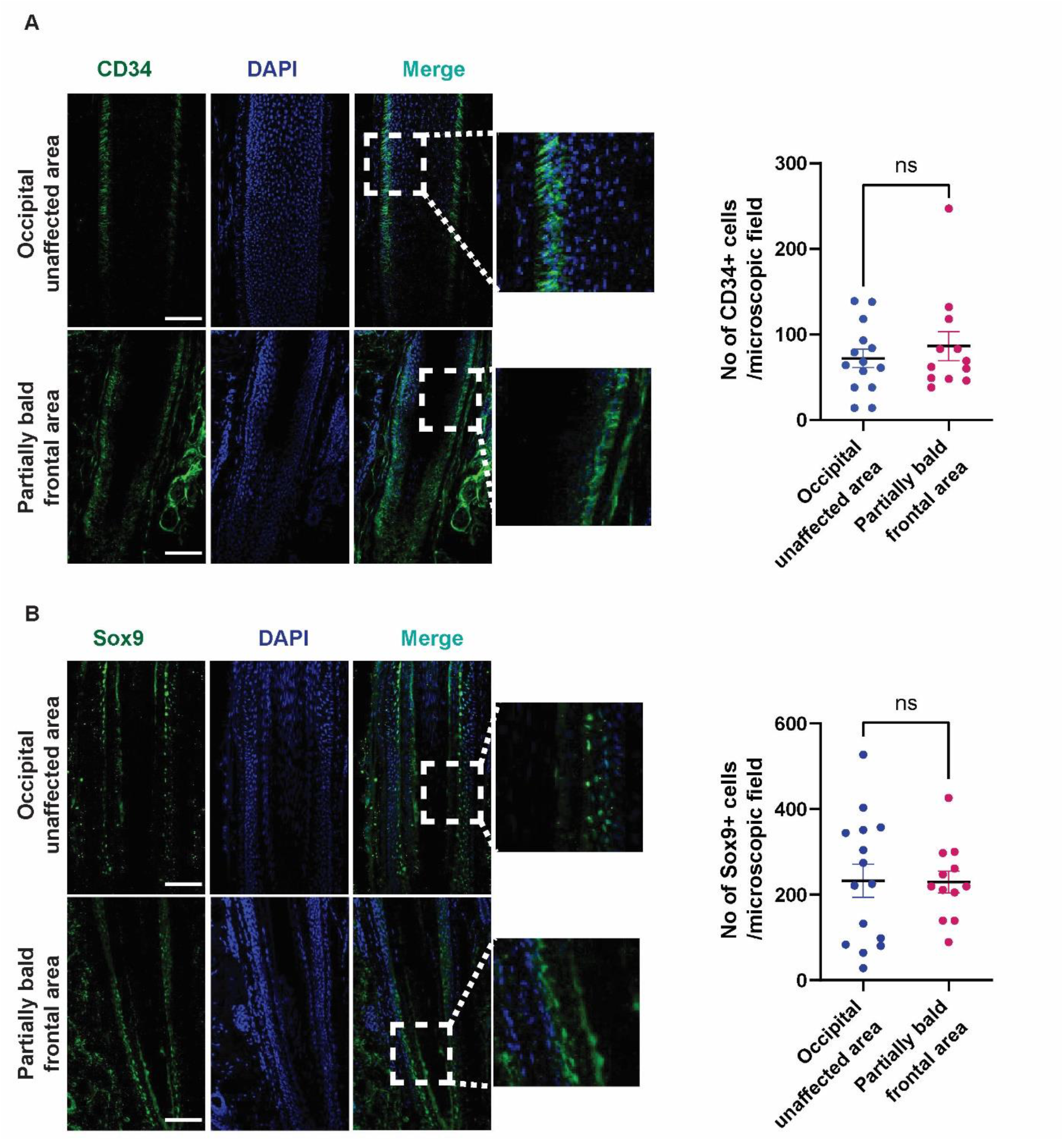
Partially bald area of AGA scalp retains CD34^+^ and Sox9^+^ progenitors. Immunofluorescence staining (green) for (A) CD34^+^ and (B) Sox9^+^ hair follicle progenitor cells in the unaffected occipital and partially bald frontal area of AGA scalp reveal that these progenitor cells are retained in the partially bald frontal area. Nuclei are marked by DAPI in blue. Scale bars: 100 µm. Quantification indicates no significant difference in number of CD34^+^ or Sox9^+^ cells between the two areas. n = 6. Data are presented as mean ± S.E.M. Each dot represents the number of progenitor cells found per microscopic field. p-values were calculated using nonparametric Wilcoxon matched-pairs signed-ranks test. p<0.05 was considered significant; “ns” indicates “not significant”. Scale bar: 100µm.

## Discussion

Recent studies have shed light on the defect in the stem cell to progenitor cell conversion in the AGA hair follicles (Garza et al., 2011), highlighting the importance of characterizing various progenitor cells in the AGA scalp. Previous works have primarily focused on Caucasian patient samples and since the prevalence of AGA differs with ethnicity, our study provides novel insights into the progenitor cell status in AGA in the Indian population. We found that the hair follicles in the completely bald area of AGA lack progenitor cells expressing CD34 and Sox9 but they are retained in the partially bald area. This indicates that progenitor cells likely diminish during the terminal stage and there is a window of opportunity for therapeutic intervention.

Androgenic alopecia is one of the most common dermatological problems worldwide for which the treatment is sought (Kaliyadan et al., 2013). The cost of 5 year treatment for AGA ranges from hundreds to thousands of dollars depending on the type of treatment (Nestor et al., 2021a). Although studies have shown that AGA can cause significant psychological distress in patients (Bade, 2016), there are limited therapeutic options available due to the lack of a complete understanding of the cellular and signaling mechanisms underlying AGA progression. Previous work have reported that androgens, especially dihydrotestosterone (DHT) dysregulate entry of hair follicles into the anagen phase by impairing HFSC differentiation via upregulating the production of DKK-1 which is an antagonist of WNT signaling (Leirós et al., 2017). Current treatment modalities often involve Finasteride which is used to inhibit the production of DHT. Minoxidil is another compound used in the treatment of AGA, which increases hair growth by shortening the telogen phase and causing premature entry of those hair follicles into anagen phase (Messenger & Rundegren, 2004). Alternative therapies include hormonal and platelet rich plasma therapies, but these have been shown to have limited effectiveness and are prone to several side effects (Mysore et al., 2019; Nestor et al., 2021a). While hair transplant is very effective in all stages of AGA, this method is costly and time consuming (Nestor et al., 2021b). Intradermal injection of autologous stem cells is an exciting new therapy (Elmaadawi et al., 2018). Progenitor cell enriched micrografts are emerging as a novel therapy (Ruiz et al., 2020), but the long term effectiveness remains to be determined. Elucidating the underlying mechanisms of AGA would help to optimize these therapeutic approaches.

Therapies aimed at activating/maintaining stem and progenitor cells would theoretically be more effective at the stage when there is initiation of active hair loss and deterioration in hair quality (partial alopecia). On the other hand, therapies intended to stimulate new hair growth in completely bald areas need to focus on replenishing the source of stem and progenitor cells.

## Materials and methods

### Human subjects

10 patients were included in the study. Diagnosis of AGA was established by the characteristic distribution of frontal and vertex hair. Patients more than 20 years old and diagnosed as male AGA, type III to VI using Hamilton –Norwood classification were included. Written informed consent for the procedure was obtained from the participants. Demographic details of all patients, including age, family history and duration of disease are summarized in Table 1. Approval to extract patient samples with informed consent was obtained from the Ethical Committee of Sapthagiri Institute of Medical Sciences and Research Centre (Bangalore, India). Analysis of patient skin samples in the Jamora lab (Bangalore, India) was approved by the Institutional Ethics Committee of the Institute for Stem Cell Science and Regenerative Medicine (Bangalore, India) (Human Ethics Approval Certificate number inStem/IEC-8/003).

**Table 1:**
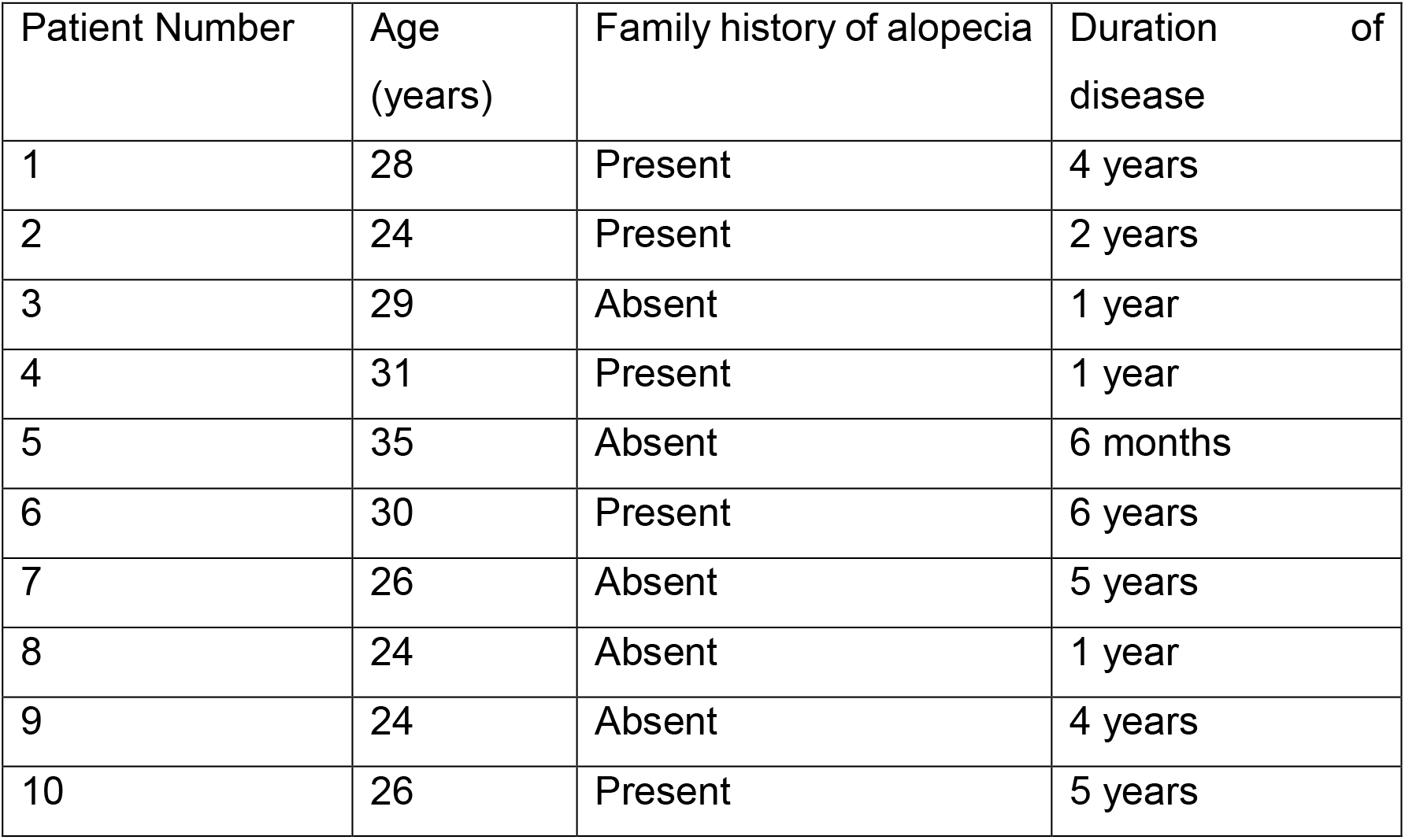
Clinical characteristics of alopecia patients.

| Patient Number | Age (years) | Family history of alopecia | Duration of disease |
| --- | --- | --- | --- |
| 1 | 28 | Present | 4 years |
| 2 | 24 | Present | 2 years |
| 3 | 29 | Absent | 1 year |
| 4 | 31 | Present | 1 year |
| 5 | 35 | Absent | 6 months |
| 6 | 30 | Present | 6 years |
| 7 | 26 | Absent | 5 years |
| 8 | 24 | Absent | 1 year |
| 9 | 24 | Absent | 4 years |
| 10 | 26 | Present | 5 years |

### Sample collection

Three mm punch biopsy specimens were obtained from all the patients from both frontal area of scalp with partial loss of hair and the occipital scalp skin that serves as a control. From 4 patients out of these, biopsies were also collected from completely bald frontal area. The samples from the 3 different areas of biopsy have been denoted as Occipital area, Partially bald frontal area, and Bald frontal area.

### Tissue preparation and histology/ immunofluorescence

Skin samples were fixed in 4% paraformaldehyde solution for 24 hours at 40C. Tissues were quenched in 125 mM glycine solution in Tris-buffered saline for 30 minutes and then incubated in 5%, 10%, and 20% sucrose solution for 6 hours, 12 hours, and 24 hours, respectively, before embedding in tissue freezing medium (Leica). Sections that were 30 µm thick were sliced using a cryostat. H&E staining was used for viewing gross follicle histology. For immunofluorescence staining, the following primary antibodies and dilutions were used: Sox9 (Abcam, catalog number ab185230, 1:200), CD34 (Abcam, catalog number ab81289, 1:100). Alexa Fluor 488 - labeled anti-rabbit secondary antibody (Jackson ImmunoResearch Laboratories) was used at a dilution of 1:200. Hoechst stain was used to mark the nuclei.

### Image collection and analysis

Imaging was performed with an Olympus IX73 microscope (Olympus), FV1000 or FV3000 confocal microscope (Olympus). Images were analyzed on the Fiji (ImageJ) software.

### Statistical analysis

Quantification of the number of Sox9^+^ and CD34^+^ cells in the occipital and frontal areas of the scalp in 6 patient sample sets was done using the Paired, non-parametric, one-tailed, Wilcoxon matched pairs signed rank test. GraphPad Prism 5.02 (GraphPad Software) was used for all statistical analyses. P < 0.05 was considered significant.

## Author contributions

Conceptualization: AH, DS, SAS, CJ; Funding Acquisition: SAS, CJ; Investigation: AH, DS, SAS; Methodology: AH, DS, SAS, CJ; Supervision: CJ; Writing – Original Draft Preparation: AH, DS, CJ; Writing - Review and Editing: AH, DS, CJ

## Acknowledgements

The authors would like to thank the Jamora lab members for their critical review of the work and insightful discussions. This work was supported by grants from the Department of Biotechnology of the Government of India (BT/PR8738/AGR/36/770/2013) and the Rajiv Gandhi University of Health Sciences (Bangalore, India) (RN003) and core funds from the Institute for Stem Cell Science and Regenerative Medicine (Bangalore, India). We thank the Central Imaging and Flow Cytometry Facility of the Bangalore Life Sciences Cluster for experimental support. The work was done in Bangalore, India.

